# Quantitative characterization of random partitioning in the evolution of plasmid-encoded traits

**DOI:** 10.1101/594879

**Authors:** Andrew D. Halleran, Emanuel Flores-Bautista, Richard M. Murray

## Abstract

Plasmids are found across bacteria, archaea, and eukaryotes and play an important role in evolution. Plasmids exist at different copy numbers, the number of copies of the plasmid per cell, ranging from a single plasmid per cell to hundreds of plasmids per cell. This feature of a copy number greater than one can lead to a population of plasmids within a single cell that are not identical clones of one another, but rather have individual mutations that make a given plasmid unique. During cell division, this population of plasmids is partitioned into the two daughter cells, resulting in a random distribution of different plasmid variants in each daughter. In this study, we use stochastic simulations to investigate how random plasmid partitioning compares to a perfect partitioning model. Our simulation results demonstrate that random plasmid partitioning accelerates mutant allele fixation when the allele is beneficial and the selection is in an additive or recessive regime where increasing the copy number of the beneficial allele results in additional benefit for the host. This effect does not depend on the size of the benefit conferred or the mutation rate, but is magnified by increasing plasmid copy number.

## Introduction

Plasmids occur naturally in bacteria, archaea, and eukaryotes [Couturier *et al.*, 1988; Wang *et al*, 2015; Meinhardt *et al*, 1990]. Most commonly found in bacteria, plasmids are typically small, extra-chromosomal, stretches of DNA that replicate independently of the host genome. Plasmid copy number, the average number of plasmids per cell, can vary from ~1 copy per cell to hundreds depending on the mechanism regulating plasmid replication. Due to their abundance, plasmids are a key source of genetic diversity and play an important role in evolution, and recent work has demonstrated the ability of high copy number plasmids to expedite acquisition of antibiotic resistance [Harrison and Brockhurst, 2012; San Millan *et al.*, 2017]. Plasmids are also widely used in synthetic biology and biotechnology, where the ability to rapidly engineer a plasmid and transform it into a host for maintenance and expression is of great value. Despite their widespread abundance in the natural world, and their frequent use in synthetic biology and biotechnology, the evolution of plasmid-encoded traits remains incompletely understood.

Plasmids have diverse inheritance modes [Novick, 1987; Summers, 1991; Williams and Thomas, 1992]. Plasmids with high copy numbers typically rely on random binomial partitioning at cell division to ensure both daughter cells receive the plasmid [Summers and Sherratt, 1984]. This strategy is not viable for plasmids with low copy numbers as there is a high likelihood a daughter cell would not inherit the plasmid [Peterson and Phillips, 2008]. Low copy number plasmids instead use active partitioning systems which distribute an equal number of plasmids to each daughter cell during division [Ogura and Hiraga, 1983; Motallebi-Veshareh *et al.*, 1990; Williams and Thomas, 1992].

Due to the independent nature of mutations, a given cell can contain a mix of plasmids where some contain the mutation and others do not. Thus, for both low and high copy number plasmids a key feature of their inheritance is the random partitioning of a potentially mixed pool of alleles into daughter cells (Figure 1).

**Figure 1:**
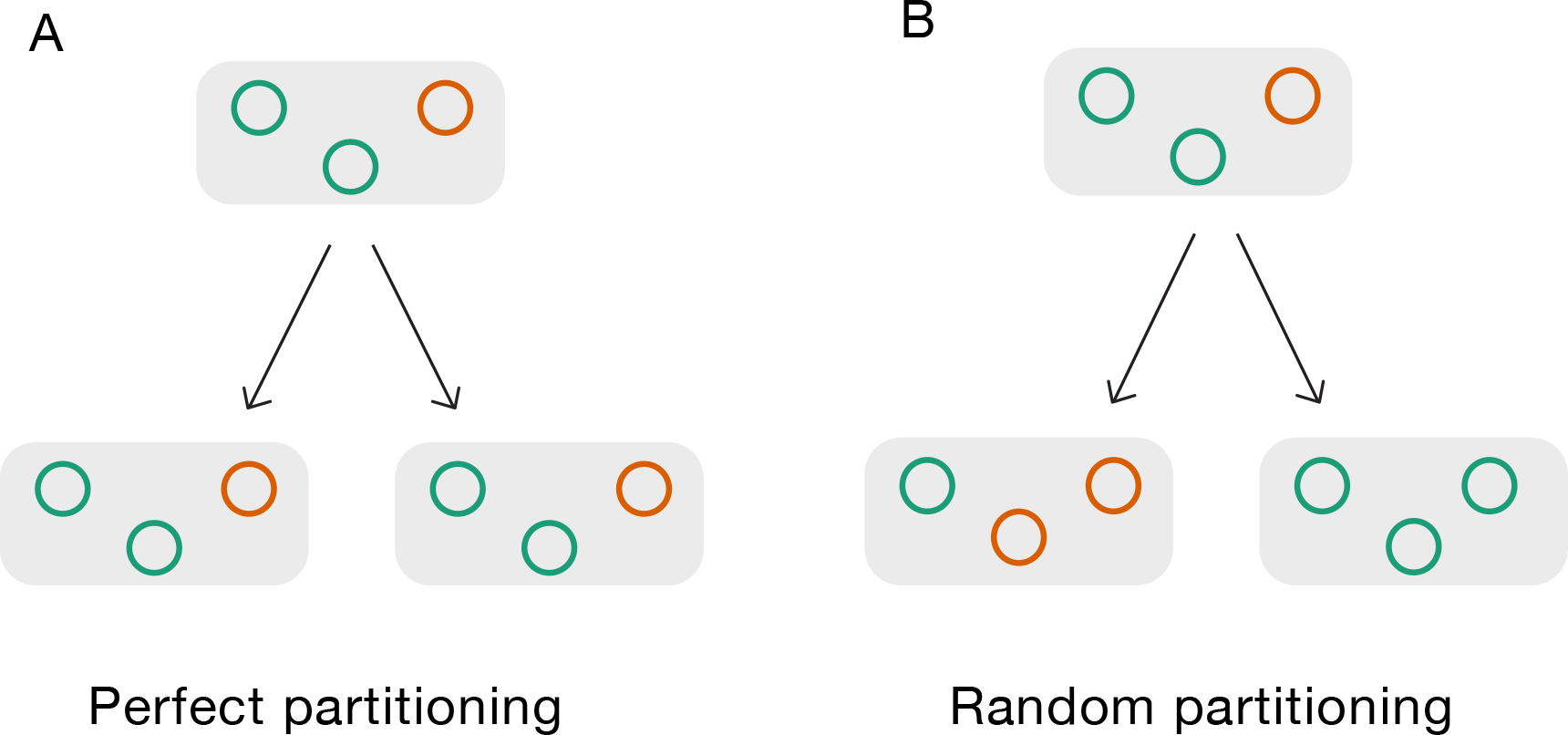
Perfect vs. random partitioning. (A) In perfect partitioning every division results in two daughter cells that both have the same distribution of functional (green) and broken (orange) plasmids. (B) In random partitioning the two daughter cells have the same total number of broken and functional plasmids as the perfect partitioning case, but the two daughter cells do not have to be identical to the parent cell or each other. In this example one daughter cell received both broken plasmids, while the other received only functional plasmids.

Previous work compared the evolutionary stability of a constitutive protein production cassette repeatedly integrated onto the genome to a similar copy number plasmid [Tyo *et al.*, 2009]. Removing the ability of the circuit to randomly partition by integrating it onto the genome dramatically increased the evolutionary stability of their protein production circuit. Recent work by Ilhan and colleagues combined experiments and simulations to understand the role random plasmid partitioning plays in the absence of selection [Ilhan *et al.*, 2018]. Their results demonstrate random partitioning increases genetic drift for mutations that have no selective advantage.

We explore the impact of random plasmid partitioning on evolvability using stochastic simulations. We find that in additive and recessive selection regimes, where having multiple copies of the mutant allele confers more benefit than the presence of a single copy, random plasmid partitioning dramatically reduces the time it takes for the advantageous allele to overtake the population. By systematically exploring parameter space, we demonstrate that this effect is independent of the total burden imposed by the plasmid and the mutation rate of the plasmid, but increases with increasing plasmid copy number.

## Results

We first consider the use case of plasmids in biotechnology applications where each copy of the plasmid encodes a function (typically production of a protein) that imposes a growth penalty on the host cell. We simulated a population where each cell contains a set of plasmids, and each functional plasmid places an independent fitness penalty on the host [Scott *et al.*, 2010]. Mutations can inactivate individual plasmids, removing the fitness penalty the plasmid introduced. All plasmids are replicated exactly once immediately before cell division and plasmid partitioning. Mimicking the active partitioning mechanisms commonly employed by low copy number plasmids, our random plasmid partitioning model distributes the same total number of plasmids to each daughter cell, but the distribution of mutant vs. functional plasmids follows a hypergeometric distribution (see “Plasmid partitioning distribution” under Methods for description). These simulations are then compared to the perfect plasmid partitioning model where each daughter cell’s plasmid contents are an identical copy of the mother cell’s plasmid distribution. The perfect plasmid partitioning case is used to represent multiple copies that are integrated onto the chromosome. This allows for independent mutation of each copy, but perfect partitioning of the group of copies into both daughter cells.

To investigate random vs. perfect partitioning, we first simulated a copy number 3 plasmid where each mutated copy of the plasmid results in a 3.3% increase in growth rate (cell with three functional plasmids growth rate = 0.9, one mutated plasmid growth rate = 0.933, two mutated plasmids growth rate = 0.966, three mutated plasmids growth rate = 1.0) both with perfect partitioning and random partitioning (Figure 2).

**Figure 2:**
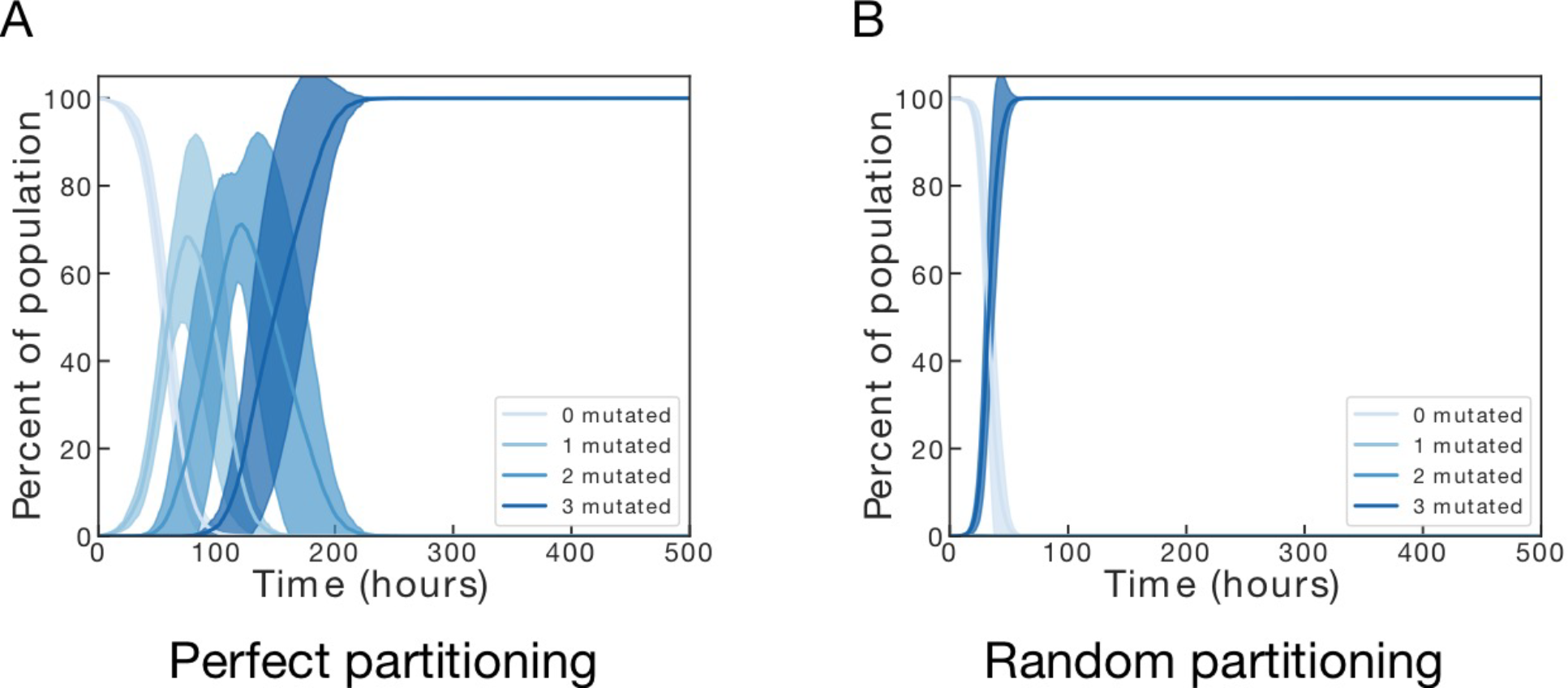
Perfect vs. random partitioning. (A) Perfect partitioning stochastic simulations. (B) Random partitioning stochastic simulations Each plot is the result of ten independent stochastic simulations. Solid lines represent the mean of the ten simulations, shaded area is +/− one standard deviation on each side. For all simulations the cell population size was capped at 10^4^, the copy number set to 3, the total burden set to 0.3, and the mutation rate set to 10^−4^.

Random plasmid partitioning not only greatly decreases the amount of time until the mutant allele becomes fixed in the population, but also forces the population to rapidly transition from the fully unbroken to fully broken state (Figure 2B).

While this result matches previously published models, it is unclear if this phenomenon is dependent on a specific set of parameters or if it is a feature common to random plasmid partitioning. This striking qualitative change in the evolutionary trajectory between perfect and random partitioning models has four key parameters that require investigation: total burden, mutation rate, plasmid copy number, and selection mode.

### Effect of random plasmid partitioning is independent of mutation rate and total burden

Two key parameters frequently considered in models of evolution are mutation rate and mutational benefit – the growth rate advantage associated with the mutation. Here, following our motivating example of a plasmid coding for constitutive expression of a burdensome protein, each mutation of an individual plasmid alleviates a fraction (1 / plasmid copy number) of the total burden. While increasing the total burden will predictably decrease the time for the mutated plasmid population to take over in both the perfect and random partitioning models, it is unclear whether the relative difference between random and perfect partitioning will be altered. To investigate this question we picked a single copy number (3) and mutation rate (1e-4) and simulated three different total burdens: 0.03, 0.1 and 0.3 both with perfect and random partitioning. As expected, increasing total burden greatly reduced times to takeover (Figure 3A, 3B). Surprisingly, changes to the burden parameter do not significantly alter the relative impact that random partitioning has on the system (3C). Random partitioning accelerates evolutionary adaptation in the additive selective regime independent of the burden the plasmid imposes on the cell.

**Figure 3:**
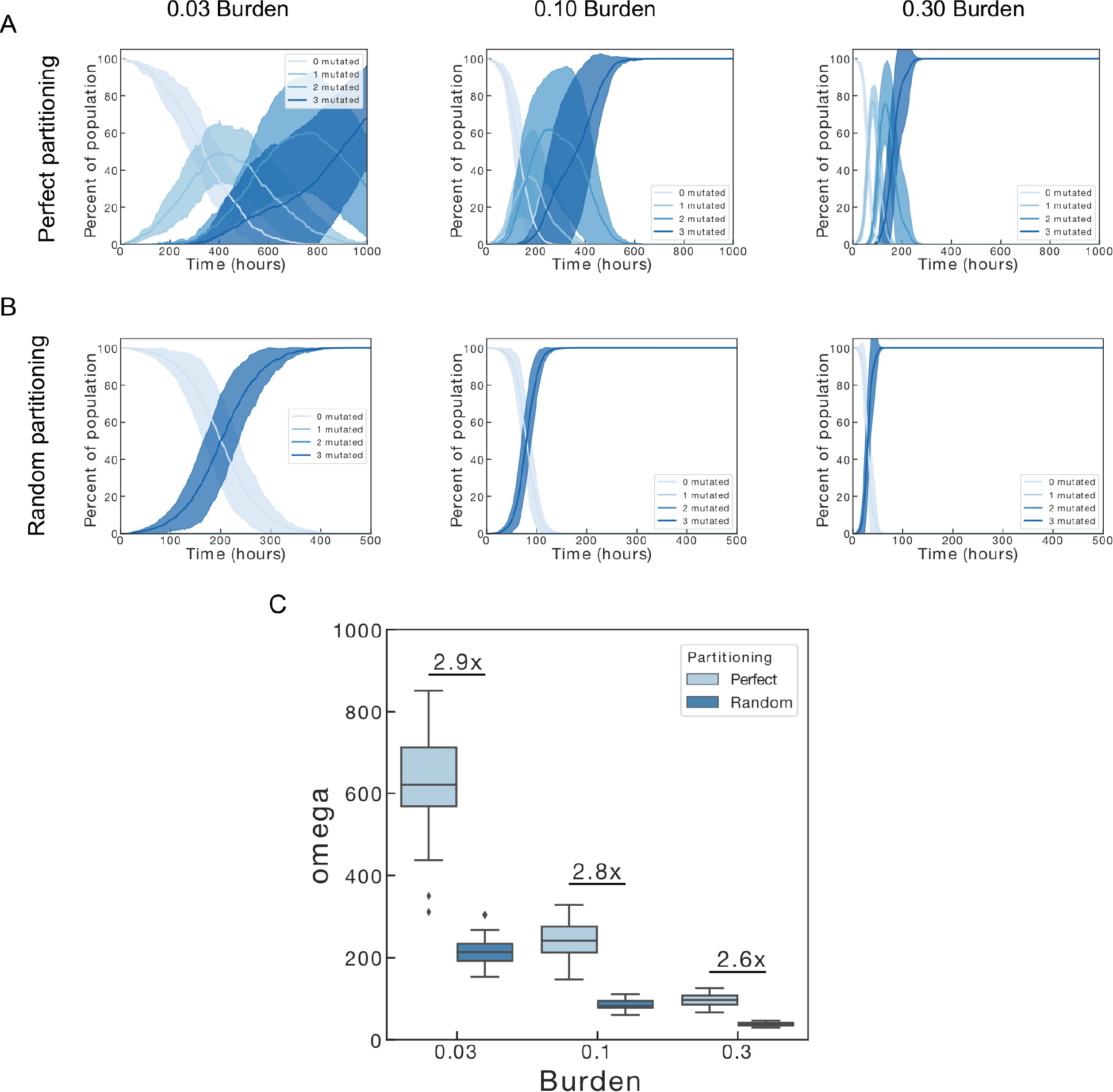
Perfect and random partitioning across increasing total burden. (A) Perfect partitioning stochastic simulations for 0.03, 0.1, and 0.3 total burden. (B) Random partitioning stochastic simulations for 0.03, 0.1, and 0.3 total burden. (C) Omega (time until half of the plasmids in the population are non-functional) as a function of mutation rate for both perfect and random partitioning simulations. Each plot in (A) and (B) is the result of ten independent stochastic simulations. Solid lines represent the mean of the ten simulations, shaded area is +/− one standard deviation on each side. The summary plot in (C) is the result of 24 independent stochastic simulations. For all simulations the cell population size was capped at 10^4^, the copy number set to 3, and the mutation rate set to 10^−4^.

The next model parameter we considered was plasmid mutation rate. Like changing total burden, changing plasmid mutation rate will change the time until mutant plasmid takeover with higher mutation rates leading to faster adaptation. However, it is difficult to intuit whether changing plasmid mutation rate will alter our finding that random plasmid partitioning accelerates evolutionary adaptation. To determine the impact of changing mutation rate on random plasmid partitioning relative to perfect plasmid partitioning, we picked a single copy number (3) and total burden (0.3) and simulated three different mutation rates, 10^−6^, 10^−5^, and 10^−4^ both with perfect and random plasmid partitioning (Figure 4A-B). As expected, increasing the mutation rate decreases both the variance in the time for mutants to dominate the populations the total amount of time it takes for them to do so. Interestingly, changing plasmid mutation rate by a factor of 100 does not significantly alter the relative difference between the perfect and random plasmid partitioning models.

**Figure 4:**
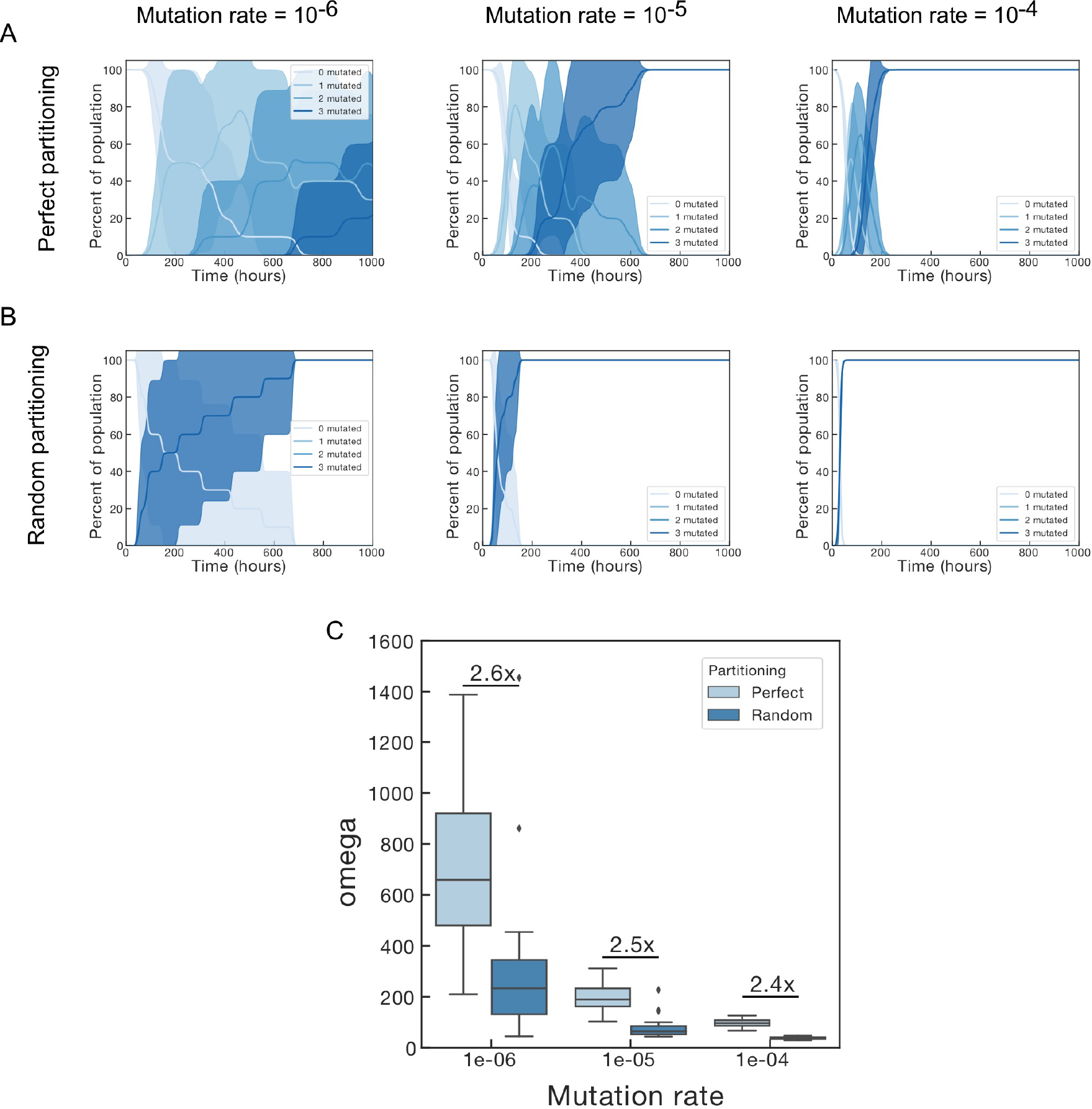
Perfect and random partitioning across increasing mutation rate. (A) Perfect partitioning stochastic simulations for 10^−6^, 10^−5^ and 10^−4^ mutation rates. (B) Random partitioning stochastic simulations for 10^−6^, 10^−5^ and 10^−4^ mutation rates. (C) Omega (time until half of the plasmids in the population are non-functional) as a function of mutation rate for both perfect and random partitioning simulations. Each plot in (A) and (B) is the result of ten independent stochastic simulations. Solid lines represent the mean of the ten simulations, shaded area is +/− one standard deviation on each side. The summary plot in (C) is the result of 24 independent stochastic simulations. For all simulations the cell population size was capped at 10^4^, the copy number set to 3, and the total burden set to 30%.

Taken together, these results suggest that random plasmid partitioning accelerates evolutionary adaptation is robust across a wide range of physiologically relevant mutation rate and burden parameter values.

### Increasing plasmid copy number increases the impact of random plasmid partitioning

Next we investigated how the effect of random plasmid partitioning would change with changing plasmid copy number. We tested three different plasmid copy numbers: 3, 5, and 10. Increasing the copy number of the plasmid while keeping the total fitness penalty to the host the same has two effects. First, it makes each mutation less valuable. If both a 10-copy number plasmid and a 5-copy number plasmid reduce host fitness by 10%, mutating one copy in the 10-copy number case only results in a 1% increase in fitness, while mutating one plasmid in the 5-copy number case results in a 2% increase in fitness. Second, it increases mutational supply. If mutations are independent, a copy number 10 plasmid is twice as likely to experience a mutation per unit time than a copy number 5 plasmid. Stochastic simulations allow us to dissect which of these contributions dominates (Figure 5).

**Figure 5:**
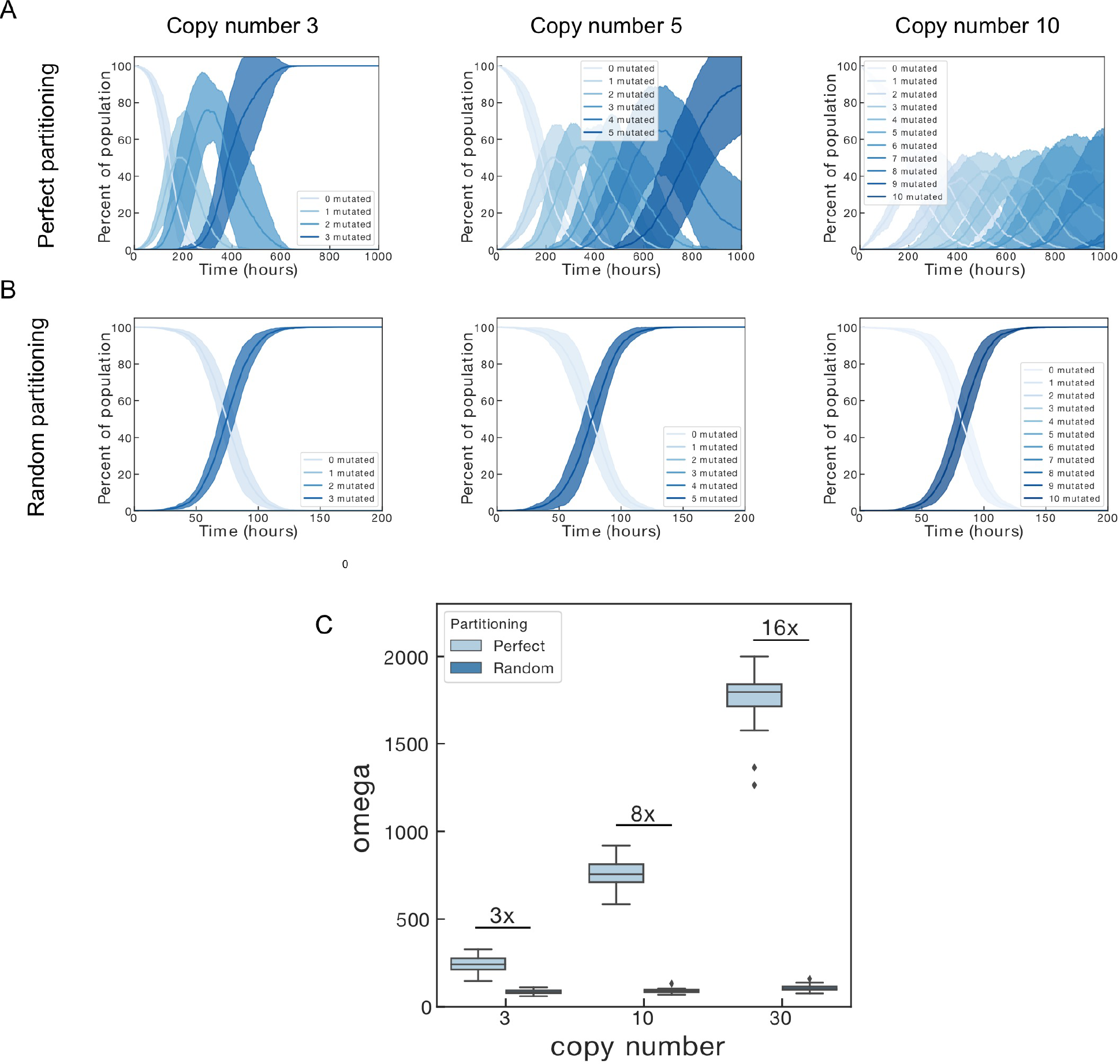
Perfect and random partitioning across increasing plasmid copy number. (A) Perfect partitioning stochastic simulations for copy number 3, 5, and 10 plasmids. (B) Random partitioning stochastic simulations for copy number 3, 5, and 10 plasmids. (C) Omega (time until half of the plasmids in the population are non-functional) as a function of copy number for both perfect and random partitioning simulations. Each plot in (A) and (B) is the result of ten independent stochastic simulations. Solid lines represent the mean of the ten simulations, shaded area is +/− one standard deviation on each side. The summary plot in (C) is the result of 24 independent stochastic simulations. For all simulations the cell population size was capped at 10^4^, the mutation rate set to 10^−4^, and the total burden set to 10%.

In the perfect partitioning case increasing copy number, while keeping the total fitness penalty constant, increases the time it takes for the mutant plasmid allele to fix in the population (Figure 5A). Surprisingly, the increase in mutational supply is more than offset by an increase in the amount of time it takes for each successive mutation (single mutant, double mutant, etc.) to become a large enough fraction of the population that an additional mutation can occur.

In the random plasmid partitioning case this effect is greatly attenuated (Figure 5B, 5C). Increasing plasmid copy number from 3 to 30 changes the takeover time by less than a factor of 2 in the random partitioning case, compared a greater than 10-fold increase in the perfect partitioning case (Figure 5C). Thus, as plasmid copy number increases the effect of random plasmid partitioning has on time to mutant allele fixation increases relative to the perfect plasmid partitioning case.

### Random plasmid partitioning accelerates evolution in additive and recessive selection regimes and slows evolution in dominant selection regimes

So far we have considered an additive selection model where each mutation is as valuable as the previous mutation and the following mutation (Figure 6A). While this is a good assumption for a simple protein expression circuit, there are other important selection modes to consider. For circuits that aim to directly regulate cell growth rate mutating any individual copy may yield a minimal increase in cell fitness, but mutating every copy of the plasmid will allow the cell to grow without any growth regulation [You *et al.*, 2004; McCardell *et al.*, 2017]. This describes a recessive selection regime, where a cell needs to mutate every copy of the allele before it receives a fitness benefit. The other selection extreme is dominant selection where a single mutant copy conveys all the fitness benefit to the cell.

**Figure 6:**
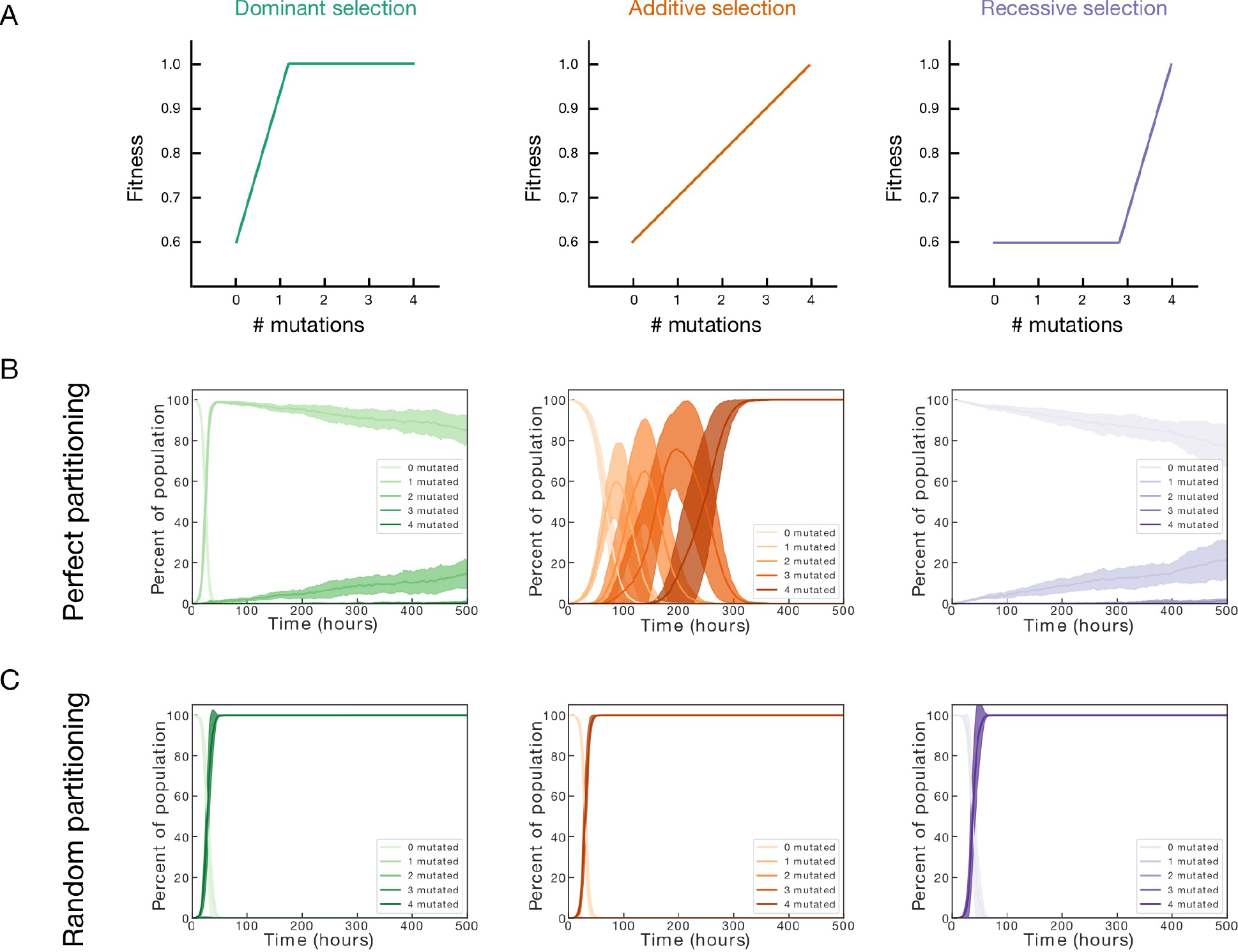
Perfect and random plasmid partitioning under different selection modes. (A) Schematic depicting cell fitness as a function of the number of mutations. In dominant selection only the first mutation is beneficial. In additive selection each mutation is equally beneficial. In recessive selection only the last mutation is beneficial. (B) Perfect partitioning stochastic simulations for each selection regime. (C) Random partitioning stochastic simulations for each selection regime. For all plots the cell population size was capped at 10^4^, the mutation rate set to 10^−4^, the plasmid copy number set to 4, and the total burden set to 30%. Each plot has ten separate stochastic simulations. Solid lines represent the mean of the ten simulations, shaded area is +/− one standard deviation on each side.

Matching previous findings, stochastic simulations reveal random plasmid partitioning slows adaptation in the dominant selection region compared to perfect plasmid partitioning [Ilhan *et al.*, 2018]. This is caused by partitioning events that generate one daughter cell without a mutant plasmid when the parent had one, unnecessarily concentrating the mutant alleles into fewer cells. This concentration can only slow evolutionary adaptation, and increase genetic drift, because all that is required for the maximum growth rate is a single mutated plasmid. In the recessive regime, however, we find that random plasmid partitioning accelerates adaptation more than in the additive regime. Without random plasmid partitioning, a given cell needs to acquire multiple independent mutations before receiving a fitness advantage, an exceptionally rare event. With random plasmid partitioning, within a few divisions one daughter cell will accumulate all the mutated plasmids, rapidly acquiring the fitness advantage associated with having all mutated plasmids. These results demonstrate that for both additive and recessive selection regimes random plasmid partitioning accelerates evolutionary adaptation, but in dominant selection regimes random partitioning slows adaptation.

## Discussion

Despite the widespread abundance of plasmids, and their importance in both natural systems and synthetic biology, the impact that random partitioning has on the evolution of plasmid-encoded traits has been underexplored. Experiments comparing many genome integrations to a similar copy number plasmid confirmed modeling predictions that random plasmid partitioning should reduce evolutionary stability for a mutation governed by additive selection [Tyo *et al.*, 2009]. A recent study by San Millan and colleagues investigated evolution of resistance to beta-lactam class antibiotics also suggested a possible role for random plasmid partitioning in accelerating adaptation [San Millan *et al.*, 2017]. Simulations and experiments by Dagan and colleagues showed that in the absence of selection, random plasmid partitioning slows evolution, despite increasing mutational supply, by enhancing genetic drift [Ilhan *et al.*, 2018].

Our simulation results demonstrate that random plasmid partitioning accelerates mutant allele fixation when the allele is beneficial and the selection is in an additive or recessive regime where increasing the copy number of the beneficial allele results in additional benefit for the host. This effect does not depend on the size of the benefit conferred or the mutation rate, but is magnified by increasing plasmid copy number.

Additional work that adds more mechanistic detail to the model, including considerations for differential plasmid replication and alternate mechanisms for plasmid partitioning, would be of interest. Further, modeling plasmids that lack active partitioning mechanisms, such as the high copy plasmid ColE1, by using a binomial rather than hypergeometric model, would be of importance due to the widespread use of high copy number plasmids in biotechnology.

Previous studies examining plasmid stability in synthetic systems largely ignore the fact that the traits are encoded on multicopy plasmids [Sleight and Sauro, 2013; Rugbjerg *et al.*, 2018]. Revisiting these studies and considering plasmid-level contributions to population-level loss of function would be of value.

Finally, our work suggests a simple strategy for increasing the stability of synthetic circuits that have additive or recessive selection. By integrating onto the genome multiple times, the circuit stability is augmented both by an increased copy number and by avoiding random partitioning upon cell division. While multiple systems already exist for integrating the same construct into the genome multiple times they have not been widely adopted [Tyo *et al.*, 2009; Gu *et al.*, 2015]. A simpler method that results in a single-step integration into multiple genomic locations would be a valuable contribution to the field of synthetic biology.

## Methods

### Simulations

Populations are simulated as groups of individual cells. Individual cells have a specified plasmid copy number. Individual plasmids within cells can mutate following the specified mutation rate. When cells divide the daughter cells are either copies of the parent cell (perfect plasmid partitioning) or undergo hypergeometric plasmid partitioning at cell division. To determine the number of broken vs. unbroken plasmids the two daughter cells get the following procedure is performed. First, the parent cell duplicates all plasmids. Then, each daughter randomly samples half of the total number of plasmids without replacement. This results in a hypergeometric distribution of broken plasmids received.

The population is simulated using the Gillespie stochastic simulation algorithm [Gillespie, 1977]. All code is written in Python3.5 using Just-In-Time compiled numba code to speed up computation [Jones *et al.*, 2014; Van Der Walt *et al.*, 2011; Lam *et al.*, 2015; McKinney, 2011; Hunter, 2007]. All code is publicly available and can be accessed at https://github.com/andyhalleran/plasmid_partitioning/tree/master/code.

Some of the simulations (specifically the higher plasmid copy number simulations) can take a few minutes to run per trajectory. To save readers unnecessary time re-running existing simulations, all summary statistics for the simulations performed for this paper can also be found at https://github.com/andyhalleran/plasmid_partitioning/tree/master/simulations. Analysis code for generating summary plots can be found at https://github.com/andyhalleran/plasmid_partitioning/tree/master/code.

### Plasmid partitioning distribution

While we believe the hypergeometric partitioning model is a fairly accurate approximation for actively partitioning plasmids, there are other distributions one could pick to model the partitioning process. Another possible plasmid distribution to consider is the binomial distribution. We did not use the binomial distribution for two reasons. First, we thought it was a poorer match to the active partitioning mechanisms that most low copy number plasmids employ. Second, the binomial plasmid partitioning mechanism can result in cells receiving 0 plasmids or gradually gaining more and more plasmids, approaching an infinite number of plasmids. To prevent these scenarios an additional mechanism driving the plasmid copy number to the desired steady state must be implemented.

## Acknowledgments

This work was funded by Caltech SURF, NIH T32 to ADH, and NSF GRFP to ADH. Research supported by the Air Force Office of Scientific Research, grant number FA9550-14-1-0060. We thank Justin Bois for providing code for the core Gillespie stochastic simulation algorithm used. The authors would also like to thank Sam Clamons, John Marken, Andrey Shur, and Rory Williams for helpful discussions.

